# A Multiscale computational model of YAP signaling in epithelial fingering behaviour

**DOI:** 10.1101/2021.12.13.472497

**Authors:** Naba Mukhtar, Eric N Cytrynbaum, Leah Edelstein-Keshet

**Affiliations:** Department of Mathematics, UBC

## Abstract

In epithelial-mesenchymal transition (EMT), cells organized into sheets break away and become motile mesenchymal cells. EMT plays a crucial role in wound healing, embryonic development, and cancer metastasis. Intracellular signaling in response to mechanical, topographic, or chemical stimuli can promote EMT. We present a multiscale model for EMT downstream of the protein YAP, which suppresses the cell-cell adhesion protein E-cadherin and activates the GTPase Rac1 that enhances cell migration. We first propose an ODE model for YAP/Rac1/E-cadherin interactions. The model dynamics are bistable, accounting for motile loose cells as for adherent slower cells. We implement this model in a cellular Potts model simulation of 2D wound-healing using the open source platform Morpheus. We show that, under suitable stimuli (depicting topographic cues) the sheet exhibits finger-like projections and EMT. Morphological, as well as quantitative differences in YAP levels as well as cell speed across the sheet are consistent with preexisting experimental observations of epithelial sheets grown on topographic features in vitro. The simulation is also consistent with experiments that knockdown or over-express YAP, inhibit Rac1, or block E-cadherin.

**SIGNIFICANCE:** In normal wound-healing, cell in an epithelium divide, grow, and migrate so as to seal a gap. In some pathological states, epithelial-mesenchymal transition (EMT) can lead to abnormal morphology, including fingering, breakage of single cells or multicellular clusters from the sheet edge. The mechanochemical control of this behaviour by cell signaling circuits (YAP, Rac1, and E-cadherin) reveals how the competition between cell adhesion and cell migration contributes to the process. We use the open-source computational platform Morpheus to investigate a multiscale model for the interactions of the proteins inside cells and the resulting morphology of the cell sheet. Results are consistent with experimental results in the literature.

## INTRODUCTION

Epithelial-mesenchymal transition (EMT) plays an important role in normal embryonic development and wound healing, and in pathology such as cancer metastasis (1, 2). In EMT, relatively static well-organized cells in epithelia lose adhesion, separate from the sheet and migrate away individually or in clusters. The migration of single cells has been studied experimentally and computationally for decades, and, as a result, there is much information about the subcellular mechanisms that regulate and tune cell motility (3–7).

In collective cell migration, cells move in structures such as swarms, clusters, or sheets. Cell group properties emerge, such as polarization, morphology, or self organization (8, 9). Various intracellular signaling networks regulate the cell-cell adhesion and cell motility that are responsible for EMT. Here we focus mainly on responses of cells to topographic features on the size-scale of the extracellular matrix (ECM). The ECM, a scaffold on which cells migrate, provides topographic cues that are detected by cell-surface (integrin) receptors, stimulating a cascade of signaling. It was shown by Park et al (10), that under appropriate conditions, such cues can lead to YAP (Yes-associated protein) signaling that promotes fingering and epithelial-mesenchymal transition (EMT) in a 2D wound-healing experiment. Park et al. hypothesized that their observations could be explained by a relatively minimal signaling circuit, consisting of YAP, the small GTPase Rac1, and the E-cadherin adhesion protein. Our motivation here is to investigate these hypotheses by constructing a multiscale model of their experimental setup and investigating sheet morphology and other quantitative features that the model predicts.

In normal wound-healing of an epithelial sheet, the expansion of the sheet front progresses uniformly to fill the wound. In various abnormal states, including invasive cancer, the sheet morphology becomes ragged, forming finger-like protrusions or losing cells and cell clusters in EMT. Park et al (10) demonstrated that the topography of the substrate (an array of nano-scale ridges mimicking the extracellular matrix) has a major effect on the shape and migration speed of the expanding front resulting in complete or partial EMT. They hypothesized that this response can be attributed to the interactions of YAP with effectors that suppress E-cadherin expression, and regulators that activate Rac1, Figure 1a.

**Figure 1:**
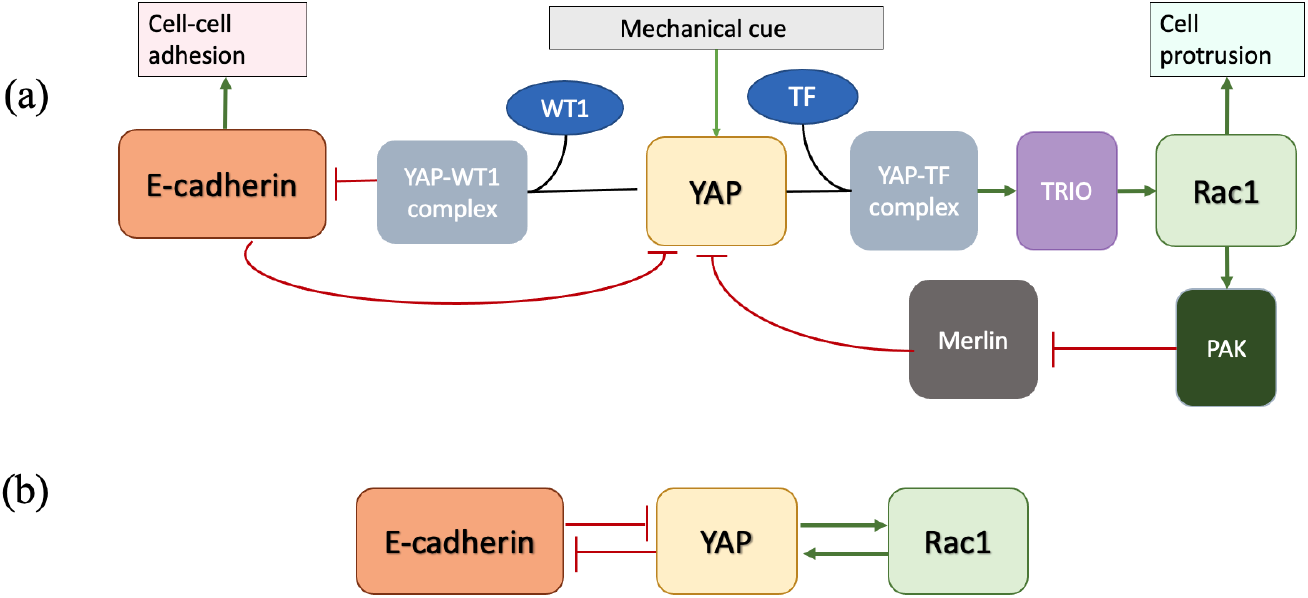
The YAP signaling system based on (10): YAP, Rac, and E-cadherin interactions with (a) detail of relevant intermediates and downstream effects on cell protrusion and adhesion. (TF= transcription factor, WT1= Wilms tumor-1, TRIO is a Rac1 GEF, i.e. activates Rac1, PAK=p21 activated kinase.) (b) The simplified system we used to represent the YAP/E-cadherin mutual antagonism and YAP-Rac mutual positive feedback.

YAP is activated by mechanical or topographic cues such as ECM fibers or nano-ridge arrays (NRAs) on a similar size-scale. Active YAP localizes to cell nuclei, where it binds to Wilms Tumor-1 (WT1), and acts as a transcription factor that suppresses E-cadherin expression. High YAP activity hence leads to loss of cell-cell adhesion. At the same time, results in Park et al (10) suggest that E-cadherin inhibits YAP activation and nuclear localization, so that YAP and E-cad are in a mutual negative feedback loop.

YAP interacts with the GTPase Rac1, and thereby affects cell migration. Park et al showed that YAP enhances the Rac1 GTP-exchange factor (GEF) TRIO, that then activates Rac1. Signaling from Rac1 via PAK and Merlin then also promotes YAP activation, indicating that YAP and Rac1 participate in mutually positive feedback. These interactions suggest that high YAP cells should have low adhesion and high motility, whereas low YAP cells should be relatively static and strongly adherent to one another.

We focus on several questions that arise from (10), specifically: (1) How does ECM signaling affect epithelial sheet morphology to promote EMT? (2) Can reduction of cell-cell adhesion through E-cadherin inhibition and increase of cell speed through Rac1 activation lead to fingering and cell dissemination in epithelial cell sheets? (3) What specific inputs from ECM to the YAP signaling networks could account for the experimental observations? Park et al (10) found that there was a sharp change in YAP activity between the front and rear of the sheet when grown on NRAs. This leads to the question: (4) What internal signaling dynamics account for this observation? (5) Are simulations of the knock-down and over-expression of YAP, E-cadherin, or Rac1 consistent with experimental manipulations of this type?

To approach these questions, we first revise an ordinary differential equation (ODE) model from (10) describing the dynamics of YAP, E-cadherin, and Rac1 (Figure 1b). We analyze the model to determine a suitable parameter regime that yields bistable YAP activity, consistent with observations in (10). In the second step, we create a multicellular multiscale simulation where the YAP intracellular signaling is implemented in each cell, while minimal additional assumptions are made for cell-cell adhesion, communication, and cell motility. The model Rac1 level is linked to cell migration speed, and the model E-cadherin level sets the cell-cell adhesion in the simulated cell sheet.

There are many multicellular-migration computational models in the recent literature (see (11) for a recent review). In (12, 13), vertex-based models are used to simulate wound healing. In (14), a Voronoi polygon model is used with multiple cell phenotypes to obtain finger-like projections. Monolayers in which cells are spheres or solid objects include (15), where the main focus was on adhesion between two distinct cell types. The role of cell-cell junctions in sheet expansion was explored in (16).

A proliferation of disparate computational models for collective cell migration has appeared in recent literature. Some of these simulations also include intracellular signaling. However, synthesizing an overall understanding of the results faces the challenge that computational details, decisions made in modeling, parameter values, and other conditions are often missing or hard to reproduce. Even where simulation code is shared, obstacles due to compatibility and technical issues remain.

To overcome such problems, we used the open-source software Morpheus (17), as it makes multiscale modelling both accessible, and easy to share. The same platform can be used to build up ODE models for the internal cell circuit, to represent cell shapes, migration, and cell-cell interactions, and to visualize the collective behaviour of a group of cells. Our results are linked to small (.xml) files that allow readers to exactly replicate the behaviour we report, and capture all parameters, conditions, and assumptions used. Furthermore, as we demonstrate later on in this paper, the same framework can be easily adapted to study wound-healing governed by other signaling systems, such as TGF*β*.

## METHODS

### The YAP-Rac-Ecadherin model

To study the dynamics of YAP (*Y*), Rac (*R*), and E-cadherin (*E*), we revised a model from the Supplementary Information of (10). In that work, two models were described, one for YAP-Rac, and a second for YAP-E-cadherin. Since all three interact in the cell, we combined these into a single model. Furthermore, we revised the original model equations of (10) in several ways, partly to avoid negative concentrations, and partly to accommodate the constant total pools of Rac and YAP in the cell. See (18) for details.

YAP and Rac are activated and inactivated: *C* ⇆ *C*_*I*_, where *C* and *C*_*I*_ are the active and inactive forms, respectively. The equations for the activated forms have the structure

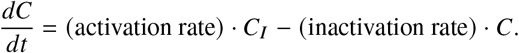

We assumed that synthesis and degradation of both YAP and Rac1 are negligible on the time scale of interest so that YAP and Rac1 have roughly constant total amounts (*Y*_tot_, *R*_tot_) in each cell:

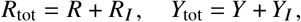

Hence, the inactive forms, *R*_*I*_ and *Y*_*I*_, can be eliminated. We propose the following system of equations for YAP, Rac1 and E-cadherin.

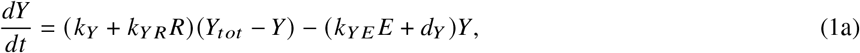

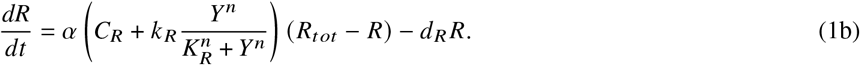

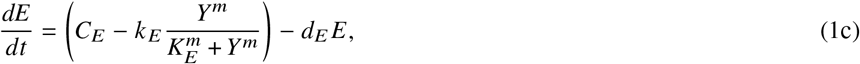

In these equations, *k*_*Y*_ and *C*_*R*_ are basal rates of activation of, respectively, YAP and Rac1, and *α* = 1 for control cells. *C*_*E*_ is the basal E-cadherin production rate. The parameters *k*_*YR*_ and *k*_*R*_ are magnitudes of feedback-associated enhancement of activation (of YAP and Rac) while *k*_*E*_ is the YAP-associated damping of the E-cadherin synthesis rate. We assume that *C*_*E*_ *> k*_*E*_ to ensure that the E-cadherin synthesis rate is non-negative. The parameters *d*_*Y*_, *d*_*R*_ are rates of inactivation of YAP and Rac, and *d*_*E*_ is the E-cadherin rate of turnover. Finally, *k*_*YE*_ is the rate of E-cadherin-mediated YAP inactivation.

Key experiments in (10) included YAP knockdown (KD) and YAP overexpression (OE). We modeled these manipulations by lowering (KD) or raising (OE) the total YAP pool, *Y*_*tot*_. To mimic the experiments in which the Rac1-GEF TRIO was inhibited, we take 0 ≤ *α* < 1.

### Differential equation solver

The YAP/Rac/E-cadherin equations were solved numerically in XPPAUTO (19), and bifurcation plots were created using the in-built AUTO feature. The same equations were then coded in Morpheus as a preliminary step. (See Supplementary Information Figure 2.)

### Morpheus simulations

Morpheus (17) is a freely available, multiscale, modelling platform and simulation environment with a convenient graphical user interface and shareable xml files. We first assembled the ODE model for intracellular YAP signaling (SI Figure 2). We then embedded the YAP model within individual “cells”, represented by Cellular-Potts Model option of Morpheus. Details of settings used in Morpheus are described further on, in the Supplementary Information, and in (18).

### Multiscale model implementation

The following assumptions were made in assembling the morphology model:

1. Intracellular signaling components are assumed to be well-mixed inside each CPM cell.
2. The nano-ridge array (10), not explicitly modeled, is represented as a directional cue. The directed motion of cells is assumed to be in the *x* axis direction.
3. Cell speed is influenced by Rac1 activity according to a directional force with strength given by

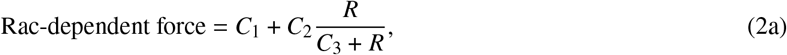

where *C*_1_ is basal migratory speed, *C*_2_ is an increase in speed due to active Rac1, and *C*_3_ is the Rac1 level that results in speed increase by 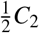.
4. Cell-cell adhesion depends on E-cadherin according to

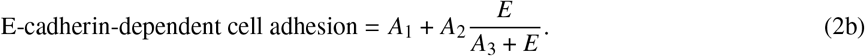

We generally took *A*_1_ = 0, and implemented Morpheus AddOn adhesion by the above formula. Ecad-independent adhesion is set by a default cell-cell contact energy parameter in the CPM simulations (see SI).
5. Only cells at the sheet edge are “induced” to respond to NRAs, and do so by an elevated Rac1 activation rate, *C*_*R*_.
6. The influence of induced cells is assumed to spread by elevating the parameter *C*_*R*_ locally in neighboring cells. At each time step, a cell’s *C*_*R*_ value has some small probability of being augmented by a small fraction of the average *C*_*R*_ value in its neighbourhood, up to some maximal *C*_*R*_ value.

We ran tests with single cells and cell pairs to optimize the adhesion and cell speed for various cell states. See Supplementary Information and SI Figure 3.

### Boundary conditions and initialization

We simulated the cell sheet on a rectangular domain. Potts model cells are initialized at the left boundary of a 2D domain, and assumed to divide with some probability near that edge. The left boundary is sealed. The domain is assumed to be periodic in the y direction. Cells that arrive at the right edge “flow past” and are removed from the simulations.

The basal Rac1 activation, *C*_*R*_, is initially set to a low value (*C*_*R*_ = 0.001) in each cell. To mimic the NRA mechanical stimulus that results in YAP activation, a small number of cells at the sheet edge are randomly endowed with a higher *C*_*R*_ value, chosen to place the cell state in the basin of attraction of the high-YAP steady state (Figure 2 blue bifurcation curve).

**Figure 2:**
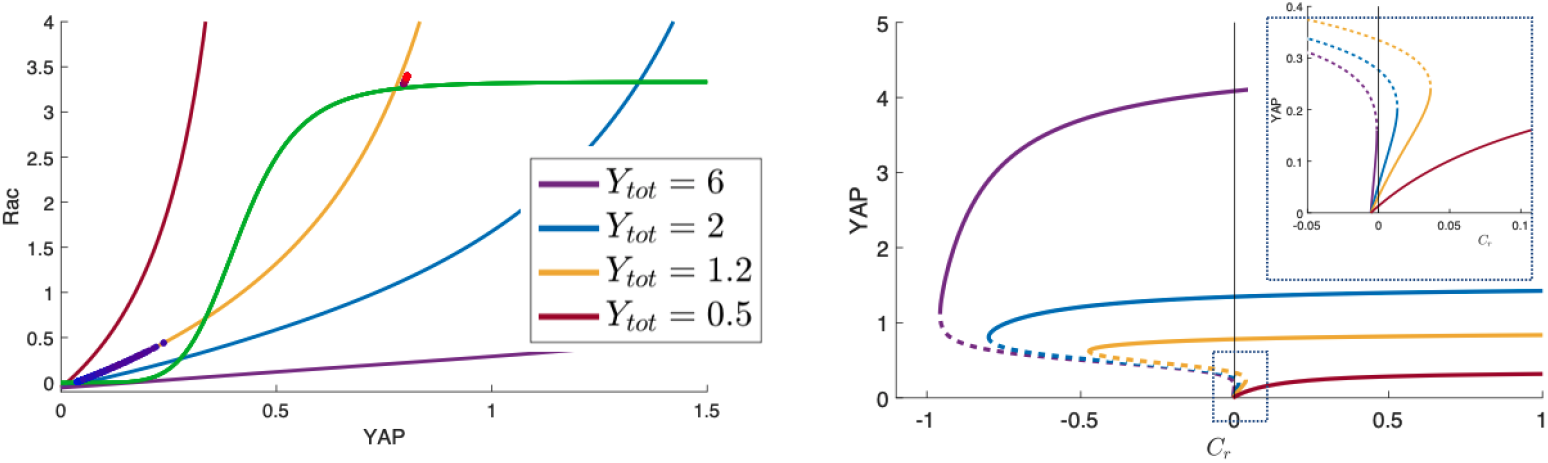
Left: A projection of the YRE model (1) onto the YAP-Rac plane, showing the YAP nullcline for control cells (*Y*_*tot*_ = 1.2, yellow), YAP knockdown (*Y*_*tot*_ = 0.5, red), YAP over-expression (*Y*_*tot*_ = 2, blue) and a higher, monostable, level (*Y*_*tot*_ = 6, purple). Steady states are at the intersections of green and red/yellow/blue/purple curves. The (YAP,Rac1) concentrations in individual cells from the control Morpheus simulation shown in SI Fig. 5 are plotted as dots, coloured from blue to red corresponding to the cells’ position in the domain (leftmost blue, rightmost red). Note that the cell states cluster around the two stable steady states with trailing cells (left, blue) being at low YAP/Rac1 and those on the leading edge of the sheet (right, red) being at high YAP/Rac1. Right: Bifurcation diagram for the same three *Y*_*tot*_ values, with *C*_*R*_ as the bifurcation parameter. For YAP KD (red), active YAP remains low and only increases slightly with *C*_*R*_. For Control,(yellow): there is a regime of bistability. YAP OE, (blue): YAP is high for all *C*_*R*_ values except for a narrow interval of bistability near *C*_*r*_ = 0. The inset shows a zoom of the region near *C*_*r*_ = 0. Parameter values as in Table 1. Produced by the XPP file: YapRacEcad.ode

## RESULTS

### Bistability in the YRE equations

Plotting the intersections of the YAP and Rac null-surfaces (*Y*^′^ = 0, *R*^′^ = 0) with the E-cadherin null-surface (*E*^′^ = 0) and projecting those onto the YAP-Rac plane, we found that the number of steady states varied with parameter values, as shown in Figure 2. For values of *Y*_*tot*_ *<* 0.86, there is a single stable steady state with low YAP/Rac and high E-cadherin. For *Y*_*tot*_ *>* 5, there is also a single stable steady state, but with high YAP/Rac and low E-cadherin. Between these *Y*_*tot*_ values, the system is bistable. For *Y*_*tot*_ ≈ 2, the system is bistable but the high YAP high Rac state dominates, in the sense that most initial conditions fall into its basin of attraction.

We treat the value *Y*_*tot*_ = 1.2 as representative of the control experiment scenario, with KD and OE values being *Y*_*tot*_ = 0.5 and *Y*_*tot*_ = 2. The KD value is monostable. While the OE value is bistable, the upper steady state tends to dominate, as we will subsequently show. In SI Figure 2, we show the time-dependent solutions *Y*(*t*), *R*(*t*), *E*(*t*) of Eqs. (1). For the control parameter values, the model is bistable, and initial conditions determine whether a high YAP or a low YAP state emerges. For YAP KD (*Y*_*tot*_ = 0.5) only one low YAP state remains. For YAP OE (*Y*_*tot*_ = 2), most initial conditions evolve to the high-YAP steady state.

The same conclusion can be seen from the bifurcation diagrams (Right panel, Figure 2). Here we use the basal rate of Rac1 activation, *C*_*R*_ as bifurcation parameter, since it later represents a property that varies across a cell sheet. We find that control cells (yellow curve, Fig. 2) can have high YAP wherever *C*_*R*_ is sufficiently high, e.g. at the sheet front, and a regime of bistability elsewhere behind the leading edge of the sheet.

### Cell sheet simulations

We asked whether the YAP signaling model, together with basic further assumptions about cell-cell interactions would recapitulate experimental observations of (10). To investigate this question, we implemented a cell sheet simulation. Each cell has the same internal YAP-Rac1-Ecad signaling model, and each cell has its individual state and trajectory.

Several additional assumptions were required to scale up from one cell to many, and to assemble the Morpheus simulations, as described in Methods. For example, since active Rac1 promotes F-actin assembly (not explicitly modeled) and cell protrusion, so we assumed that higher Rac1 activity leads to faster cell migration. Similarly, the level of E-cadherin was linked to cell-cell adhesion. Additionally, while the NRAs may stimulate all cells, experimental evidence demonstrates that only cells close to the leading sheet edge respond with nuclear (active) YAP. Hence, we assumed that initially, only cells at the sheet edge (not inhibited by contacts with neighbors) would be “induced” to respond with an elevated Rac1 activation rate, *C*_*R*_. As leading cells migrate outwards at the edge of a cell sheet, they create tension on cells behind them. In biological cells, this tension inactivates the protein Merlin, abrogating Merlin’s inhibition of Rac1. In the computations, we do not model Merlin explicitly, but we assume that the rate of Rac1 activation (*C*_*R*_) can locally spread, as described in Methods.

Parameters and rates of processes are not easily measurable in experiments, so we tuned parameters of the system to correspond with YAP bistability in the control cultures, and with the overall KD and OE results, as shown in Figure 2. Before finalizing the full wound-healing computation, we ran test with single cells and cell pairs to optimize the adhesion and cell speed for various cell states. Such tests are described in the Supplementary Information and shown in SI Figure 3. We discuss parameter values in greater detail further on.

#### Formation of fingering, control simulation

The above YAP/Rac1/E-cadherin model, computational steps, and initial conditions were implemented. A time-sequence of the resultant sheet morphology is shown in Figure 3.

**Figure 3:**
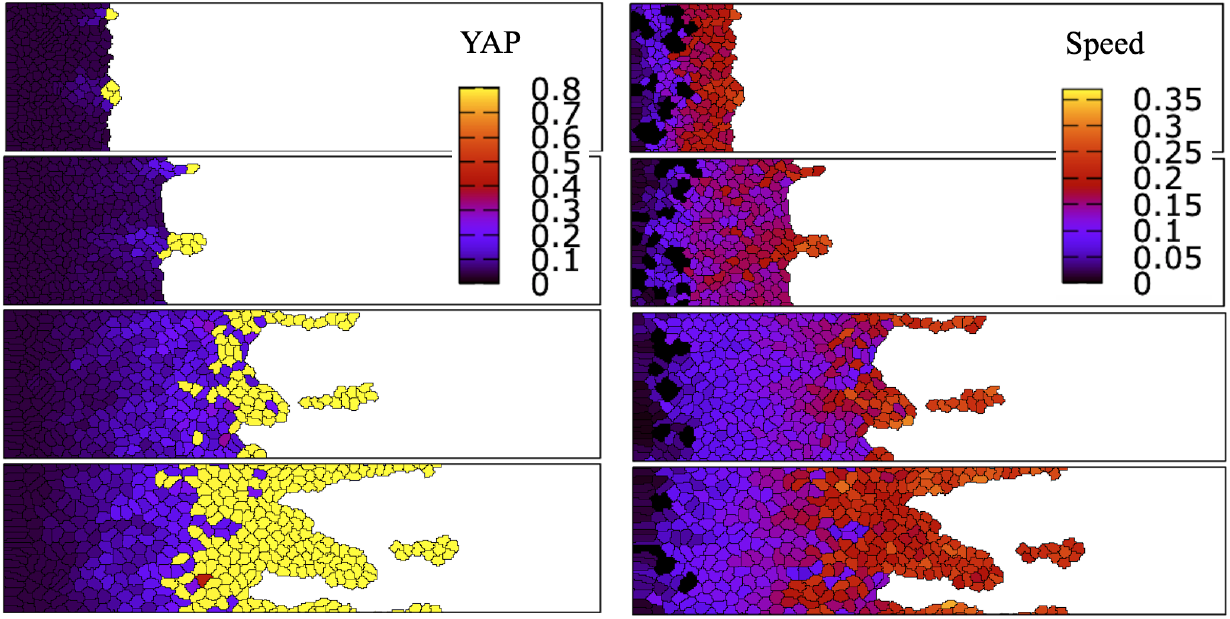
Formation of fingers in the multiscale simulation of a control cell sheet on topographic (NRA) substrate. A time sequence (top to bottom, *t* = 250, 500, 1000, 1250) showing sheet morphology colored for YAP and cell speed. Cells near the front of the sheet have high YAP. In the middle of the sheet there is a mixture of cell states. Morphology is comparable to Figure 2c of (10). Parameter values as in Table 1. Morpheus File: YAPspeed.xml. A movie is [here](https://www.dropbox.com/s/kd93wdjo70exyz7/YAPspeed.avi?dl=0).

At several places along the sheet edge, we find small bumps that grow into fingers, with occasional dissemination of small clusters. The fingers are relatively straight and rarely merge. In Figure 3, we show the YAP level (left) and cell speed (right). The same time sequence with levels of Rac1 and E-cadherin is displayed in SI Figure 5. Cells near the front of the sheet have high YAP and Rac1 activities, and low E-cadherin, while cells near the back have low YAP, low Rac1 activity, and high E-cadherin. The middle of the sheet consists of both cell types. A few high-YAP cells in the middle of the sheet originate from spreading influence (*C*_*R*_) of the leaders at the edge. Only some of them are sufficiently induced through this influence to transit to the high-YAP steady state.

The simulation morphology results are qualitatively consistent with experimental results shown in Figure 2c of (10). We also find that cell speed increases with proximity to the sheet edge, as observed in the experiments.

#### Parameter values

We comment here on the tuning of parameters that resulted in the fingering shown in Figure 3. The values of the Morpheus parameters governing inter-cellular adhesion and motility, specifically *C*_1_, *C*_2_, *A*_2_, and cell-cell and cell-medium contact energies, were appropriately fine-tuned. In particular, the cell speed values *C*_1_ and *C*_2_ had to be large enough for collective migration to be driven primarily by cell motion rather than pressure of cell division at the back of the sheet. At the same time, it had to be low enough to prevent the sheet from breaking apart prematurely. The adhesion parameters (*A*_2_) and the cell-cell and cell-medium contact energies were set so that inter-cellular adhesion would maintain a cohesive sheet, apart from occasional small dissemination of a cluster or two. At the same time, it was set to avoid overpowering cell migration altogether. In general, we found that morphology was highly sensitive to the balance between cell division, cell removal, and cell speed. Many other morphologies are possible when such parameters are modified.

#### Cell trajectories

We compared cell behaviour for cells on NRA versus cells on flat substrates. For growth on a flat substrate, we simply assumed that no cells get induced with high Rac1 activity at the leading edge of the sheet. We found that the sheet edge remained smooth and no fingering occurred. (See SI Figure 4 for sheet morphology on a flat substrate.)

Observing the actual cell trajectories and speeds in the simulation, we found behaviour that was consistent with experimental observations of (10) in both the flat and NRA wound-healing cases, as shown in Figure 4.

**Figure 4:**
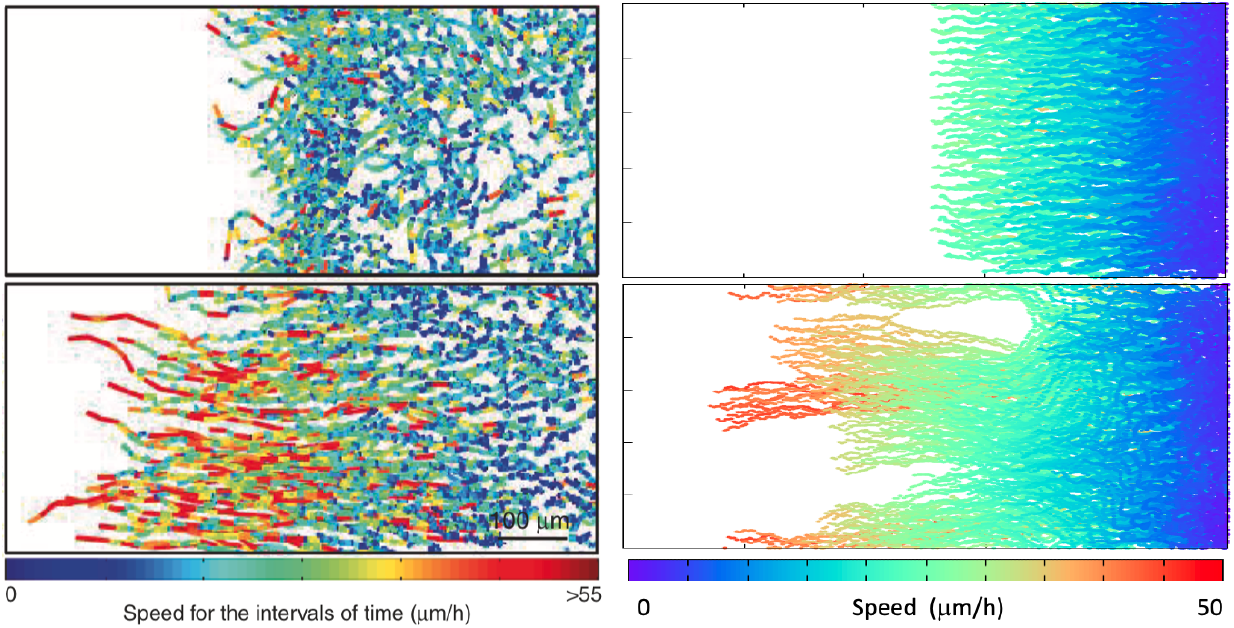
Trajectories for cell sheets grown on flat (top) and NRA (bottom) substrates. Left: Experimental results from Fig 1b of Park et al (10) (licensed under a Creative Commons Attribution 4.0 license). *t* = 6 hours, speed measured every 20 minutes). Right: Simulations, with speed scaled to experimentally observed range. Cell speeds are relatively slow and uniform on a flat substrate, compared to NRA substrate, where some cells at the leading edge are moving faster. (Simulation shown for t = 1800, in a domain of size 100× 500 *μ*m). Parameter values as in Table 1. Morpheus simulation files: top, flat substrate - CellTrajCntrFlat.xml and Movie, bottom, NRA substrate - CellTrajCntrNRA.xml and [Movie](https://www.dropbox.com/s/wqfdr9y10tkgw65/CellTrajCntr_NRA.avi?dl=0).

### YAP knockdown and over-expression

Experiments in (10) were designed to investigate the effects of knockdown (KD) and overexpression (OE) of YAP levels in the cell sheet. As previously noted, we represent KD (OE) by increasing (decreasing) the total YAP, *Y*_tot_. We ran the same simulation as before, and compared the dynamics and morphology.

Morphological results are provided in SI Figures 6 and 7. Briefly, YAP KD simulation (SI Fig. 6) consists of cells with very low active YAP throughout the sheet, and slow sheet expansion. In the YAP OE simulation (SI Fig. 7), multiple fingers form and grow early on, and cells rapidly jump to the high-YAP state with only a small increase in *C*_*R*_. These observations are consistent with experiments in (10). However, in our YAP KD simulations, fingers are wider than those seen in YAP KD experiments of (10).

We can understand some of the results from the underlying cell model. For high *Y*_*tot*_ (YAP OE) the is high-YAP steady state dominates for most cells in the range of values of the bifurcation parameter *C*_*R*_. This implies that YAP activity will be high throughout most of the sheet. For low *Y*_*tot*_ (YAP KD) there is a single low-YAP steady state, so YAP is low across the sheet. For moderate *Y*_*tot*_ (control cells), there is a transition from low YAP (at the back) to high YAP (at the front), since the inducing stimulus (value of *C*_*R*_) is graded from front to back. See bifurcation results in Figure 2, and note that the value of *C*_*R*_ is correlated with distance from the front of the sheet. For the control simulation, the level of active YAP vs distance from front edge is displayed in the right panel of Figure 5. We see high YAP at the front, and low YAP at the back, with a zone of mixed high/low YAP cells in the middle of the sheet. Such results are consistent with experimental findings in Park et al. (10) Fig 7a.

**Figure 5:**
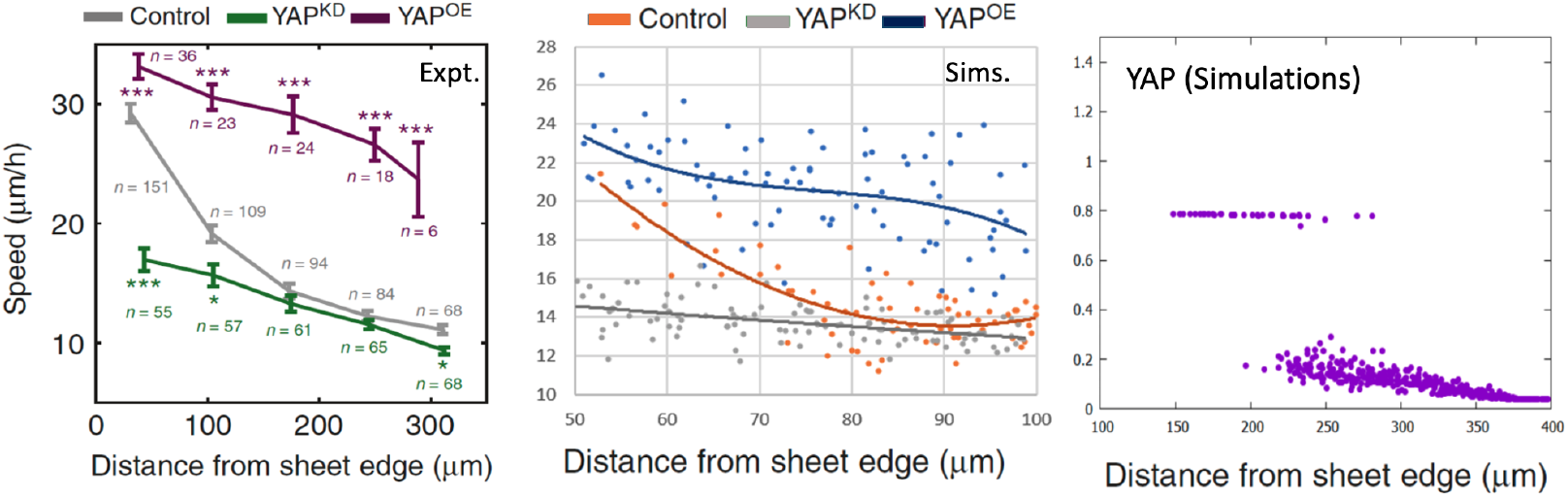
Distribution of YAP and cell speeds across the migrating sheet. Right: In the control NRA simulation, YAP activity is highest at the front of the sheet, and low in the back. Left: Experimental cell speed vs distance for YAP knockdown (KD), control, and overexpression (OE) from Figure 2b of Park et al (10), (licensed under a Creative Commons Attribution 4.0 license.) Center: Simulation results. *Y*_*tot*_ = 0.5, 1.2, 2 for KD, control, and OE respectively; other parameter values as in Table 1. (*t* = 1000) Morpheus file: nra.xml

### YAP affects migration speed across the sheet

We asked whether the differences in speeds observed in experimental YAP manipulations in (10) agree quantitatively with our computational results. Consequently, we plotted cell speed versus distance from the leading edge of the cell sheet. Results are shown in Figure 5. As shown in the left panel (data from (10)), speed is always higher at the leading edge. However, the gradient of the speed is relatively shallow in both high and low YAP conditions, but steep in the control case. Simulations (central panel of Figure 5) are consistent with these observations. We can understand this behaviour from the dynamics of the YAP-Rac1-Ecad model, since the control is in a bistable regime where the front edge and rear of the sheet are at distinct steady states. In the KD YAP, the single steady state changes gradually with distance from the front edge, but does not have a sharp jump. In YAP OE, most of the sheet is in the high YAP state, so again a sharp jump is not evident. The relatively good agreement of simulation and experimental quantitative results provides a measure of confidence in the ability of the minimal signaling model to depict the behaviour of the wound-healing experiments.

#### E-cadherin blocking and Rac1 inhibition

In (10), antibodies were used to block the adhesive function of E-cadherin. These experiments led to less stable fingers and more breakage of cells and clusters compared to control cell sheets. We simulated this experiment by reducing *A*_2_, the E-cadherin dependent part of the cell-cell adhesion. The YAP-Rac-Ecad model and parameters were kept as before. The results of this change compared to control cells are shown in the left panels of Figure 6, where we also compare the number of disseminations in control versus 50% E-cadherin function blocking (FB). We found that fingers in the E-cadherin blocked sheet morphology are quite thin compared to the control sheet. The number of disseminations also increased significantly (bar graph).

**Figure 6:**
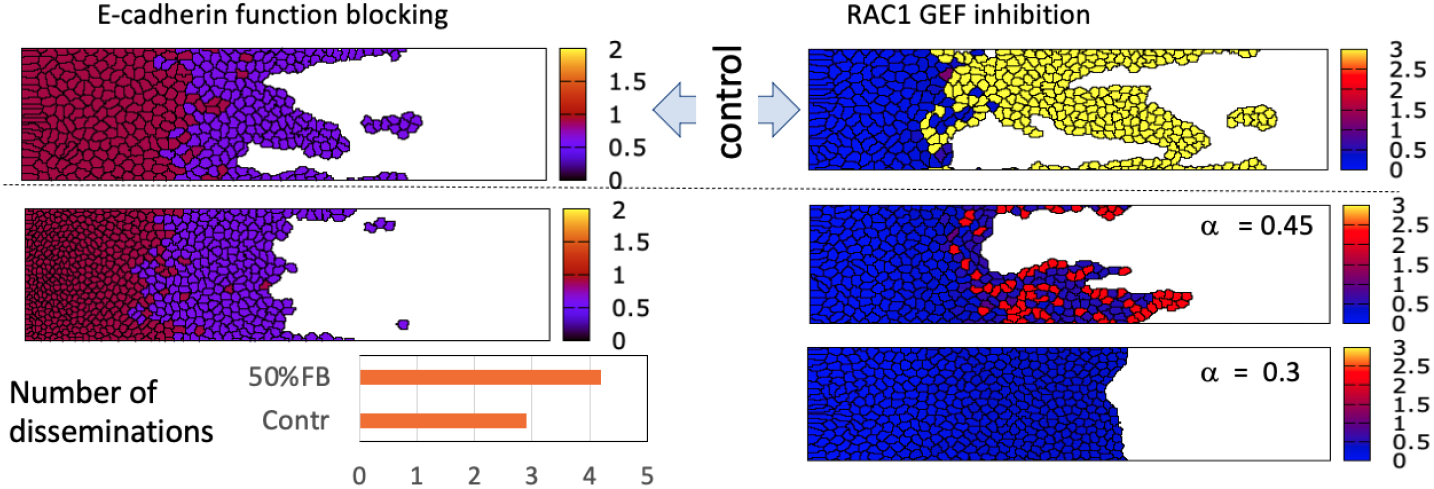
E-cadherin function blocking (FB) and Rac1 inhibition. Left: reducing the cell-cell E-cadherin adhesion (governed by the parameter *A*_2_) leads to thinner fingers and more breakage of cells (image at *t* = 1250). Right: reducing the Rac1 activation rate (*α* < 1) in (1b) slows down the sheet expansion and eventually abolishes fingering (images at *t* = 1500). In each case, the control is shown as the top panel. Color map shows E-cadherin levels (left) or Rac1 levels (right).

In (10), Rac1 inhibition was experimentally imposed with an inhibitor (NSC 23766) that binds to the Rac GEF TRIO, preventing it from activating Rac1. To simulate this experiment, we merely reduced the Rac1 activation rate factor *α*. For control cells, *α* = 1, and for Rac1 inhibition, *α* < 1. Results of this inhibition are shown in right panels of Figure 6. Experimental Rac1 inhibition yields a sheet with fewer fingers and slower growth.The value *α* ≈ 0.45 leads results that are consistent with experimental observations in (10). For strong inhibition, (*α* = 0.3) fingering no longer takes place, a prediction that could be tested experimentally.

## DISCUSSION

In this paper, we linked the dynamics of YAP, Rac1, and E-cadherin in individual cells to intercellular adhesion and cell migration in an epithelium. In so doing, we could investigate effects on morphology, including the formation of finger-like projections. We could also quantify the distribution of cell states and cell speeds across the sheet, as well as cell dissemination from the sheet edge.

To do so, we adapted a preexisting model from the Park et al. (10) SI, and used the publicly available platform, Morpheus, and its multiscale CPM capabilities. The cell-cell adhesion and migratory capabilities were then dictated by internal cell states as well as by the neighborhood of the cells. We obtained qualitative and quantitative agreement with experimental observations of (10) both in control sheet growth on a topographic substrate with nano ridge arrays (NRAs) and with various experimental manipulations of YAP, Rac1, and E-cadherin.

To arrive at consistent behaviour, we had to tune parameters of the ODE model (placing the model in a regime of bistability for the control NRA case) and the parameters of cell-cell and cell-NRA interactions in the multiscale Morpheus simulation. We had to assume, for example, that a small number of cells at the front of the sheet were stimulated by the topography, while cells at the back, where contact inhibition is greatest, were less responsive initially. While we assumed that induced cells initially had high Rac1 activation rate *C*_*R*_, this assumption is to some extent arbitrary: any parameter value that promotes high YAP in the given cell would similarly lead to a transition to the high-YAP cell state. Another requirement was that cell-cell communication would spread the high rate of Rac1 activation (*C*_*R*_) from induced cells to their neighbors. Other biological factors (discussed below) may be at play in real cells.

In (10), it was found that in epithelia grown on NRAs, YAP activity switches from a high level at the front to a low level at the rear of the sheet. We showed that, given an appropriate choice of parameters, a bifurcation in the ODE model for the signaling circuit explains the observation. (See blue bifurcation curve in Figure 2 and right panel of Figure 5.) The rate of Rac1 activation, *C*_*R*_ is correlated with distance from sheet edge, since it is elevated at the sheet front and spreads rearwards. This explains the experimental controls. In YAP KD and YAP OE, (adjusting only one model parameter for the total YAP in cells, *Y*_*tot*_), there is no longer bistability, so YAP distribution as well as cell speed increases much more gradually towards the sheet front.

Our predictions agree with experiments in which YAP or Rac1 or E-cadherin were manipulated in several ways. E-cadherin function-blocking resulted in increased breakage of cells from the sheet, Rac1 knockdown yielded a sheet with slow finger growth, and YAP over-expression and knockdown resulted in high and low cell speeds throughout the sheet respectively. In the latter, proximity to the sheet edge had little effect on speed compared to the control case. These predictions are consistent with the experimental counterparts in (10). We also found some dissimilarities. For example, in YAP KD, a large, wide finger emerges in our simulation, as opposed to narrow fingers seen in experiments. Additionally, while the cell speed curves for simulated YAP KD, OE and control are qualitatively similar to experimental results, the absolute quantitative values were somewhat lower.

Based on results, we can address several questions raised in the Introduction. (1) ECM signaling topography, by elevating the activity of the mechanosensitive protein YAP, can lead to cell states with higher speed and lower adhesion, that promotes EMT. (2) We found that high YAP states, with reduced cell-cell adhesion and increased cell speed can lead to fingering and cell dissemination, based on our multi scale simulations. (3) ECM induction of high YAP signaling either directly, or by pathways that increase Rac activation could be the inducing stimulus that leads to fingering morphology. This is evident from the mutual positive feedback between YAP and Rac1. (4) The sharp switch in YAP activity between front and rear of NRA-grown cell sheet can be explained by virtue of the bifurcation in model for the signaling circuit. The front and rear of the sheet have cells in distinct steady states of the YAP signaling system. (5) The over-expression and knock-downs of YAP, and inhibition of E-cadherin, and Rac1 all have qualitative effects in the model that are consistent with those in the experiments.

Our computational model has several limitations. First, we implemented cell division only at the back of the sheet to allow for sheet expansion. In reality, the probability of cell division, as well as cell death is likely nonzero everywhere in a cell sheet. Cell division and death throughout the sheet could be easily implemented in a future version of the Morpheus simulation.

We considered a minimal model in which YAP partners affect adhesion and migration of cells. Other intermediates involved in the YAP/E-cadherin and YAP/Rac1 feedback loops were not included. For example, Merlin is loosely implemented in the simulation by the spread of high *C*_*R*_ (Rac1 activation rate) from cell to cell. This simplifies the actual biology. The protein Merlin is inactivated by the tension exerted on a cell by its moving neighbour, thereby increasing the basal activation rate of Rac1. Therefore in future versions of the model, Merlin could be included explicitly or spread of *C*_*R*_ could be directly linked to some measure of tension on a given cell.

The nano-ridge topography of (10) is indirectly modeled in two ways: (1) It is associated in our simulations with cell induction (high *C*_*R*_ values) at the sheet front edge and (2) it defines a directional cue for migration. In chemically-induced sheet expansion, such stimuli could stem from a morphogen or growth factor gradient (e.g. TGF-*β*). Furthermore, it would be trivial to assign the induction parameter to some other aspect of the YAP circuit such as elevated YAP rate of activation, *k*_*Y*_, for example. The model could be adapted to any other circuit controlling cell adhesion and migration, such as the beta-catenin signaling. See, e.g. (20) for self-organization of multicellular clusters based on an in-silico delta-notch-E cadherin computational study.

Movies in (10) demonstrate fluid-like sheet expansion with cells that slide past one another. In our simulations, the sheets are relatively “rigid” and cells hardly overtake one another. This fluid vs solid behaviour is likely due to CPM parameter settings that set the rigidity and stochastic fluctuations of cell shapes. Finally, as of this writing, software packages such as Morpheus cannot solve reaction-diffusion equations inside deforming domains. For this reason, cells were assumed to each be well-mixed, even though the activities of Rac1 and YAP are known to be spatially distributed in real cells.

Comparing our computations with those in the literature, there are points of similarity and some divergence. For example, while (12), used vertex-based model rather than a CPM model to describe an epithelium, they too implemented an energy-based method with target cell area and perimeter, as well as cell-cell contact energies. In (15), two cell types competed in a cell-center based model for an epithelial sheet. This study was based on the explicit forces between cells, rather than an energy minimization method. The adhesion values were preassigned, not a result of internal signaling. In the paper by (21), on the other hand, the interactions of E-cadherin and beta-catenin were incorporated into a cell-center based multiscale computation. The authors examines cells popping out of a 2D cell sheet, rather than fingering morphologies.

Alternate gene networks that regulate EMT have been considered (11). In (22) a larger beta-catenin signaling circuit is modeled using ODEs that include Rac1 and E-cadherin, as does our model, but no multiscale tissue computation. In (23), a CPM model is coupled to a finite-element model to describe EMT in a multiscale model. This model combines intra and intercellular signaling to account for EMT in wound healing and other geometries. A circuit studied in (24) explores the effect of TGF*β* on a ZEB-SNAIL circuit governing EMT. This paper, like (25), focuses on models for the time dynamics of genes, miRNAs, and proteins in the circuit, but not collective cell interactions nor morphology. However, adopting such alternate circuits to test in our multiscale simulation is relatively easy.

In recent years, there has been increased interest in cell states that are in an intermediate EM form, neither fully epithelial nor fully mesenchymal, but somewhere along the continuum (24–27). It has been suggested that this partial EM state may be more efficient at generating tumors (27). Such ideas raise the question of whether a given cell circuit that governs EMT has more than two steady states (e.g. three states, representing cell states E, EM, and M.) The simplified circuit that we studied here (Figure 1b, with assumptions that led to the system of equations (1) is bistable, but not tristable as is. In the context of experiments of (10), tristability was not needed to account for observations. However, real cells clearly have our circuit embedded inside a significantly complex signaling circuit, with multiple additional positive and negative feedback loops. It would be relatively straightforward to revise our model by adding feedbacks, and cascades or multimerization (Hill functions with powers *n* ≥ 2) to arrive at an alternative tristable version of the model, see for example (28, 29) for an indication of how nonlinearities can achieve multistability. Here we do not go down this route. Instead, we end by demonstrating an alternate EMT circuit based on TGF*β* constructed by (24) and show its morphological predictions in the SI. In that tristable model, that we implement in the same type of cell-sheet simulation, the three cell states clearly emerge along a gradient of TGF *β* from a growing sheet. Replacing the YAP-Rac-Ecad circuit by another, more intricate or complex one is straightforward once model equations and parameter values are known.

## CONCLUSION

The mechanosensitive protein YAP, by responding to topographic cues, is able to coordinate both cell-cell adhesion (via E-cadherin) and cell migration (via the GTPase Rac1). A relatively elementary model for positive and negative feedback between YAP, Rac1 and E-cadherin accounts for experimentally observed wound-healing morphology. Distinct cell states (high YAP and fast cells vs low YAP and tightly adherent cells) are predicted by them model, and their distributions in the cell sheet agree with experiments where YAP, Rac1, or E-cadherin are manipulated. Software packages such as Morpheus, by facilitating multiscale multicellular modeling, allow us to visualize the morphology of a cell sheet, YAP associated fingering, and cell breakage. The computation is easy to share, and readily adaptable to a variety of signaling circuits that are known to affect EMT and growth of epithelia.

**Table 1:** List of model parameters and Values

| Parameter | Description | Value |
| --- | --- | --- |
| $k_Y$ | Basal Rate of YAP activation ($\mu\text{Ms}^{-1}$) | 0.1 |
| $k_{YE}$ | E-cad feedback to YAP inactivation ($\mu\text{M}^{-1}\text{s}^{-1}$) | 1.8 |
| $k_{YR}$ | Rac1 feedback to YAP activation ($\text{s}^{-1}$) | 1.8 |
| $k_E$ | YAP feedback to E-cad inactivation ($\text{s}^{-1}$) | 0.9 |
| $k_R$ | YAP feedback to Rac1 activation ($\text{s}^{-1}$) | 1 |
| $K_E$ | YAP feedback to Ecad inhibition ($\mu\text{M}$) | 1 |
| $K_R$ | YAP level for $\frac{1}{2}$-max feedback to Rac1 ($\mu\text{M}$) | 0.5 |
| $d_Y$ | YAP inactivation rate ($\text{s}^{-1}$) | 2 |
| $d_E$ | E-cad decay rate of ($\text{s}^{-1}$) | 1 |
| $d_R$ | Basal Rac1 inactivation rate ($\text{s}^{-1}$) | 0.5 |
| $C_E$ | Basal Ecad synthesis rate ($\text{s}^{-1}$) | 0.9 |
| $C_R$ | Basal Rac1 activation rate ($\text{s}^{-1}$) | 0 |
| $m$ | Ecad Hill coefficient | 3 |
| $n$ | Rac1 Hill coefficient | 6 |
| $Y_{tot}$ | Total YAP ($\mu\text{M}$) | 1.2 |
| $R_{tot}$ | Total Rac1 ($\mu\text{M}$) | 5 |

## AUTHOR CONTRIBUTIONS

NM carried out the research and wrote the MSc thesis on which this paper is based. LEK designed the research and wrote the the paper. EC participated in writing the paper and in improving the results.

## ACKNOWLEDGMENTS

We thank Lutz Brusch, Joern Strass and Andreas Deutsch for making Morpheus an open source and publicly available resource, and for help with the initial cell sheet simulation. We are grateful to Dr. Rachel Hazan for pointing us to the modeling work of Prof. Xiao-Jun Tian, and to Prof. Tian for providing equations for the model in (24) that we discuss in the SI. We thank Prof Jimmy Feng and members of the Feng-Keshet group for helpful comments and feedback. All authors were supported by Natural Sciences and Engineering Research Council (Canada).

## SUPPLEMENTARY MATERIAL

An online supplement to this article can be found by visiting BJ Online at http://www.biophysj.org.

## Notes

https://www.dropbox.com/sh/8rdl90s32bj6z4e/AAAeOaA6bJXjUdUzuXt28iLpa?dl=0

### Competing Interest Statement

The authors have declared no competing interest.

## REFERENCES

1. Lee, J. M., S. Dedhar, R. Kalluri, and E. W. Thompson, 2006. The epithelial–mesenchymal transition: new insights in signaling, development, and disease. Journal of Cell Biology 172:973–981.

2. Yang, J., and R. A. Weinberg, 2008. Epithelial-mesenchymal transition: at the crossroads of development and tumor metastasis. Developmental cell 14:818–829.

3. Mogilner, A., E. L. Barnhart, and K. Keren, 2020. Experiment, theory, and the keratocyte: An ode to a simple model for cell motility. In Seminars in cell & developmental biology. Elsevier, volume 100, 143–151.

4. Ungefroren, H., D. Witte, and H. Lehnert, 2018. The role of small GTPases of the Rho/Rac family in TGF-β-induced EMT and cell motility in cancer. Developmental dynamics 247:451–461.

5. Byrne, K. M., N. Monsefi, J. C. Dawson, A. Degasperi, J.-C. Bukowski-Wills, N. Volinsky, M. Dobrzyński, M. R. Birtwistle, M. A. Tsyganov, A. Kiyatkin, et al., 2016. Bistability in the Rac1, PAK, and RhoA signaling network drives actin cytoskeleton dynamics and cell motility switches. Cell systems 2:38–48.

6. Pollard, T. D., 2019. Cell motility and cytokinesis: From mysteries to molecular mechanisms in five decades. Annual review of cell and developmental biology 35:1–28.

7. Buttenschön, A., and L. Edelstein-Keshet, 2020. Bridging from single to collective cell migration: A review of models and links to experiments. PLOS Computational Biology 16:e1008411.

8. Friedl, P., and D. Gilmour, 2009. Collective cell migration in morphogenesis, regeneration and cancer. Nature reviews Molecular cell biology 10:445–457.

9. Rørth, P., 2009. Collective cell migration. Annual Review of Cell and Developmental 25:407–429.

10. Park, J., D.-H. Kim, S. R. Shah, H.-N. Kim, P. Kim, A. Quiñones-Hinojosa, A. Levchenko, et al., 2019. Switch-like enhancement of epithelial-mesenchymal transition by YAP through feedback regulation of WT1 and Rho-family GTPases. Nature communications 10:1–15.

11. Katebi, A., D. Ramirez, and M. Lu, 2021. Computational systems-biology approaches for modeling gene networks driving epithelial–mesenchymal transitions. Computational and Systems Oncology 1:e1021.

12. Fletcher, A. G., J. M. Osborne, P. K. Maini, and D. J. Gavaghan, 2013. Implementing vertex dynamics models of cell populations in biology within a consistent computational framework. Progress in biophysics and molecular biology 113:299–326.

13. Nagai, T., and H. Honda, 2009. Computer simulation of wound closure in epithelial tissues: Cell–basal-lamina adhesion. Physical Review E 80:061903.

14. Yang, Y., and H. Levine, 2020. Leader-cell-driven epithelial sheet fingering. Physical biology 17:046003.

15. Knutsdottir, H., 2018. The multi-levelled organization of cell migration: from individual cells to tissues. Ph.D. thesis, University of British Columbia.

16. R. Noppe, A. A. P. Roberts, A. S. Yap, G. A. Gomez, and Z. Neufeld, 2015. Modelling wound closure in an epithelial cell sheet using the cellular Potts model. Integrative Biology 7:1253–1264.

17. Starruß, J., W. De Back, L. Brusch, and A. Deutsch, 2014. Morpheus: a user-friendly modeling environment for multiscale and multicellular systems biology. Bioinformatics 30:1331–1332.

18. Mukhtar, N., 2021. A simulation of epithelial sheet growth with internal signaling. Master’s thesis, University of British Columbia.

19. Ermentrout, B., 2002. Simulating, analyzing, and animating dynamical systems: a guide to XPPAUT for researchers and students. SIAM.

20. Mulberry, N., and L. Edelstein-Keshet, 2020. Self-organized multicellular structures from simple cell signaling: a computational model. Physical Biology 17:066003.

21. Ramis-Conde, I., D. Drasdo, A. R. Anderson, and M. A. Chaplain, 2008. Modeling the influence of the E-cadherin-β-catenin pathway in cancer cell invasion: a multiscale approach. Biophysical journal 95:155–165.

22. Kumar, S., A. Das, and S. Sen, 2014. Extracellular matrix density promotes EMT by weakening cell–cell adhesions. Molecular bioSystems 10:838–850.

23. Hirway, S. U., C. A. Lemmon, and S. H. Weinberg, 2021. Multicellular Mechanochemical Hybrid Cellular Potts Model of Tissue Formation During Epithelial-Mesenchymal Transition. Computational and Systems Oncology.

24. Tian, X.-J., H. Zhang, and J. Xing, 2013. Coupled reversible and irreversible bistable switches underlying TGFβ-induced epithelial to mesenchymal transition. Biophysical journal 105:1079–1089.

25. Jolly, M. K., M. Boareto, B. Huang, D. Jia, M. Lu, E. Ben-Jacob, J. N. Onuchic, and H. Levine, 2015. Implications of the hybrid epithelial/mesenchymal phenotype in metastasis. Frontiers in oncology 5:155.

26. Bierie, B., S. E. Pierce, C. Kroeger, D. G. Stover, D. R. Pattabiraman, P. Thiru, J. L. Donaher, F. Reinhardt, C. L. Chaffer, Z. Keckesova, et al., 2017. Integrin-β4 identifies cancer stem cell-enriched populations of partially mesenchymal carcinoma cells. Proceedings of the National Academy of Sciences 114:E2337–E2346.

27. Kröger, C., A. Afeyan, J. Mraz, E. N. Eaton, F. Reinhardt, Y. L. Khodor, P. Thiru, B. Bierie, X. Ye, C. B. Burge, and R. A. Weinberg, 2019. Acquisition of a hybrid E/M state is essential for tumorigenicity of basal breast cancer cells. Proceedings of the National Academy of Sciences 116:7353–7362.

28. Macía, J., S. Widder, and R. Solé, 2009. Why are cellular switches Boolean? General conditions for multistable genetic circuits. Journal of theoretical biology 261:126–135.

29. Li, T., Y. Dong, X. Zhang, X. Ji, C. Luo, C. Lou, H. M. Zhang, and Q. Ouyang, 2018. Engineering of a genetic circuit with regulatable multistability. Integrative Biology 10:474–482.

